# Autonomous bird sound recording outperforms direct human observation: Synthesis and new evidence

**DOI:** 10.1101/117119

**Authors:** Kevin Darras, Péter Batáry, Brett Furnas, Irfan Fitriawan, Yeni Mulyani, Teja Tscharntke

## Abstract

1) Autonomous sound recording techniques have gained considerable traction in the last decade, but the question still remains whether they can replace human observation surveys to sample some animal taxa. Especially bird survey methods have been tested using classical point counts and autonomous sound recording techniques.
2) We review the latest information by comparing both survey methods' standardization, verifiability, sampling completeness, data types, compatibility, and practicality by means of a systematic review and a meta-analysis of alpha and gamma species richness levels sampled by both methods across 20 separate studies.
3) Although sound recording surveys have hitherto not enjoyed the most effective setups, they yield very similar results in terms of alpha and gamma species richness. We also reveal the crucial importance of the microphone (high signal-to-noise ratio) as the sensor that replaces human senses.
4) We discuss key differences between both methods, while richness estimates are closely related and 81% of all species were detected by both methods. Sound recording techniques provide a more powerful and promising tool to monitor birds in a standardized, verifiable, and exhaustive way against the golden standard of point counts. Advantages include the capability of sampling continuously through day or season and of difficult-to-reach regions in an autonomous way, avoidance of observer bias and human disturbance effects and higher detection probability of rare species due to extensive recordings.

## Introduction

In the face of the current threats to global biodiversity, we urgently need to devise more efficient methods to measure our vanishing, under-sampled biodiversity (Haughland et al., 2010). We need larger sampling coverage on temporal and spatial scales to detect trends across regions and with time. Material and personal resources must be deployed with maximal return on investment, and to enable international cooperation and re-use of data, we need to attain a minimal bias with standardized, comparable and repeatable sampling methods.

Vertebrates pose a particular challenge for sampling because they are considerably more mobile than plants or microorganisms, often evading detection. Most vertebrate taxa are usually surveyed by direct human observation methods (e.g. point counts and transects) because capture methods are inherently more intrusive and effort-demanding. Human observers rely on aural and visual detection to count animals and identify species, but given that some insects (e.g. cicadas and orthopterans) and most terrestrial vertebrates commonly use sound (birds, amphibians, mammals, but not reptiles), passive acoustic monitoring methods have recently gained more attention (Blumstein et al., 2011).

For birds in particular, passive acoustic sampling methods have been used extensively, and many hardware (for an overview see Merchant et al., 2015,Whytock and Christie, 2016) and software solutions have been developed (Araya-Salas and Smith-Vidaurre, 2016; Katz et al., 2016; Villanueva-Rivera and Pijanowski, 2012). However, human observation survey methods are still the standard, traditional go-to method (Bibby et al., 2000; Ralph et al., 1998). Although some research has compared acoustic methods with traditional survey methods, results are controversial as some studies showed that acoustic surveys are more effective than point counts (e.g. Haselmayer and Quinn, 2000), whereas other studies concluded the opposite (e.g. Hutto and Stutzman, 2009). Anuran surveys also follow a similar pattern in that human observation surveys are most common (Heyer et al 2016), but audio surveys with passive acoustic monitoring stations are increasingly used (Aide et al., 2013). So far however, no review has summarized these studies, and bird studies provide ample material for an interesting methodological comparison.

First we summarize the evidence of existing literature based on a systematic review including a meta-analysis, and discuss the reasons leading to the previously mentioned controversial results. Then, we present the inherent advantages of either method on the example of bird surveys, focusing on a) standardization and verifiability, b) sampling completeness, c) data type and compatibility, and d) practicality. We complement the meta-analysis and the discussion with results from our own field study to analyse the survey methods’ effect on bird detection distances and abundances.

## Methods

### Systematic review and meta-analysis

#### Data collection

We searched for scientific publications on ISI Web of Knowledge (R) with the advanced search function, covering all years and all databases (search date: 12/01/2017). We used the following search string: *TS=((bird* OR avian OR avifaun*) AND (“sound record*” OR “acoustic record*” OR “automated record*” OR “acoustic monitor*” OR “recording system*”) AND (“point count*” OR “bird count*” OR “point survey*” OR “point-count*” OR “point transect*”)).* We also searched Google scholar (search date: 12/01/2017) using the following search string: *“point count” “sound recording”,* sorted by relevance, checking all search results.

We read all titles - and abstracts when needed - to determine the relevance of each study for the systematic review. Only peer-reviewed references in English were considered. In a first step, studies that discussed and compared both acoustic and observational bird survey methods were included in our systematic review. Studies focusing on a few selected species were not considered. Full texts were retrieved and read entirely. The references within the chosen publications were further checked for additional potential studies.

In a second step, studies that additionally published data on bird species richness recorded with both methods were used in the meta-analysis. The data were extracted directly from the results in the text, or from the figures using distance measuring tools in Foxit Reader (version 8.0.2, Foxit software Inc., USA), or by extracting tables from PDF files using Tabula (https://github.com/tabulapdf/tabula). When data were not available in the publication, we contacted the main author to request them and asked whether they could also provide unpublished data sets from other studies. Publications reporting results of several sub-studies, which were either distinct in study location or methodology, were treated as independent studies. Auxiliary data such as microphone model, height, signal-to-noise ratio, number of channels, and time difference between survey methods, were also extracted or requested. Whenever possible, we computed richness numbers for unlimited and also identical detection range scenarios, as in some studies, detection ranges are lower for sound recorders. Bird detection data sets obtained from unlimited distance point counts and sound recordings yielded so-called unlimited range richness values. Bird detection data subsets from point counts and sound recordings, filtered to include only detections below a common detection distance (the sound recorders' maximal range), yielded identical range richness values. When no distance data were available to truncate the point count data sets, we still used the entire data set, yielding conservative richness estimates in favor of point counts.

#### Data analysis

We used R (version 3.2.3, R core team) for all data analysis and graphing. We used the metafor R package(Viechtbauer and others, 2010) to calculate log-transformed ratios of the richness means (alpha and gamma) of both survey methods (point counts and sound recordings) for both detection range types (identical and unlimited), hereafter called log response ratios. We used multivariate meta-analysis models (metafor package, rma.mv() function) or linear mixed-effects models (nlme package, function lme()), respectively for alpha and gamma richness. We assigned the random effect to a study ID number since some publications include several sub-studies from the same author. Studies were weighted with the number of sites in alpha richness models, and with the total survey time effort in gamma richness models. Finally, we checked the robustness of our results by running subset models excluding studies which compared richness values between methods that had different detection ranges, and for which distance data were not available to correct the bias.

For the identical range scenarios, we extracted the model's intercept as the overall log response ratio, along with its confidence intervals, across all substudies. We compared the overall log response ratio to zero, representing no richness differences between survey methods. For the unlimited range scenario, we needed to account for differences in sampling area. We hypothesized that microphone signal-to-noise ratio would ultimately determine the sound detection space size relative to the unlimited point count detection space. The microphones’ signal-to-noise ratio was included as a moderator in the alpha richness models (rma.mv() function) and as a fixed effect in the gamma richness models (lme() function). The significance (at P<0.05) of the signal-to-noise ratio predictor was assessed.

The alpha and gamma richness log response ratios were back-transformed to response ratios in percent, displayed for each study along with their overall response ratio.

### Field studies

#### Forest surveys

Twenty-eight lowland rainforest plots were visited once by different observers in the province of Jambi, Indonesia (Fig S1), from April to June 2015, during the early dry season, which corresponds to the breeding season for birds in Sumatra (Voous, 1950). Some of the forest plots had previously experienced selective logging as indicated by tree stumps, and occasional hunting and bird trapping were reported by inhabitants of the region. During the plot screening, we recorded a pure tone sequence (0.5, 2, 4, 8, 12 and 16 kHz, one second long at each step) emitted at distances of 2, 4, 8, 16 and 32 meters from the recorder, which we call the sound transmission sequence. This sequence was used to assist in bird call distance estimation.

Sound recorders (SM2Bat+ recorder fitted with one SMX-II and one SMX-US microphones) were installed one day before and programmed to start recording at sunrise. Twenty-minutes point counts were carried out between 6:00 and 10:00, one minute after arriving on the plot, to avoid disturbing secretive birds. Two survey teams surveyed one plot per day each, and each team comprised one ornithologist observer and one recordist without ornithological knowledge. The recordist notified the observer of bird calls that he did not detect and recorded all calls detected by the observer to aid in identification using a directional microphone (Sennheiser ME-66 coupled to Olympus LS-3). All detected birds were identified following MacKinnon and Phillipps (1993) and their horizontal distance was measured with laser rangefinders (Nikon Laser 100 AS and Bushnell Fusion 1 Mile). For aural detections, we estimated the perching tree position and then measured the distance to its trunk, while for visual detections, the position of the perching tree was more reliable. The number of simultaneously detected individuals and the stratum of occurrence (ground, understory, middlestory, canopy, emergent tree) was noted for all detections. The autonomous sound recordings were stopped at the end of the point count. Considering that autonomous sound recorders typically collect data without human presence and can start recording earlier in sites that are difficult to access, we used twenty minute sound recordings starting 30 minutes before the point count (except one plot with 22 minutes offset) to compare the sampling completeness of both methods more fairly and avoid disturbance from human observers.

#### Forest survey data analysis

Six months after the point counts, recordings were uploaded to a website (http://soundefforts.uni-goettingen.de) designed to store and identify the birds within. The ornithologist who carried out the point count on the same day listened to the corresponding recordings while inspecting the spectrograms. The ornithologist tagged the bird calls with the species name, number of individuals, and estimated distance. The sound transmission sequence was used as reference to support the listener in estimating distances more accurately. An additional listener without ornithological knowledge listened to the same recordings to notify the ornithologist of calls that he did not detect in the recordings.

Abundance measures were rarely reported in the literature (but see Hobson et al., 2002), so we relied on our field survey to analyse abundances between survey methods, additionally to richness. Using both point count and sound recordings data, we counted the number of distinct species (species richness) and the sum of the maximum number of simultaneously detected individuals of each species (a conservative measure of abundance like in Teuscher et al., 2015) to assess the bird sampling completeness. Species richness and abundance were compared between methods using paired Wilcoxon mean comparison tests. The significance of statistical tests was assessed at a level of *P* < 0.05.

Using the field bird survey data, we investigated the bird detection frequency of each survey method in rings of equal area but increasing distance from the observer. In point count data, we estimated the direct distance to the detected bird using the observed stratum of occurrence and the horizontal distance data. For sound recordings, we used the direct distance estimates directly as noted by the listener. We analysed the detection probability in sound recordings relative to point counts by checking whether 100 randomly selected aural observations from point counts, within a horizontal distance of 50 meters and all from one observer, also occurred in the corresponding simultaneous sound recordings. We categorized the bird detection in the audio material as heard, faint, or absent.

#### Distance estimation experiment

We used caged birds to estimate the accuracy of bird call distances obtained from sound recordings. Caged birds from two different species (*Prinia familiaris* and *Acridotheres javanicus*) were rented from local habitants and brought to a mature oil palm plantation. We installed a Song Meter (with two SMX-II microphones) on a wooden pole at a height of 1.5 m and recorded the sound at 44.1 kHz. We recorded the same sound transmission sequence as in our field study, at distances of 10, 20, 30, 40, 50, 60, and 70 m to the front of the recorder, measured with the laser range finder. We placed the bird cages on the ground and recorded bird vocalizations at different distances, randomly chosen between 2 and 80 meters. The recordings were analysed blindly by KD and the distance of each vocalization was estimated aurally while listening to the sound transmission sequence as reference. When all recordings were processed, we plotted the results with a linear regression line for each call type and bird species and calculated the correlation between the measured and estimated bird distances.

## Results

### Meta-analysis

We found 23 studies with our Web of Science search string and 129 through Google scholar. 20 were relevant, and 17 had usable data for the meta-analysis. One additional unpublished data set was collected. All relevant references are listed in <u>Table 1</u>. Almost all authors responded positively and were extremely cooperative to provide summary data or even entire data sets. However, in 5 studies detection distance data were not available to correct for the point counts' larger detection range.

Overall, alpha richness values did not differ significantly between both survey methods in the identical range scenario (Fig 1 and Table S1), neither did they in subset models. Sound recordings tended to yield higher alpha richness values when compared to point counts, but lower gamma species richness.

**Figure 1:**
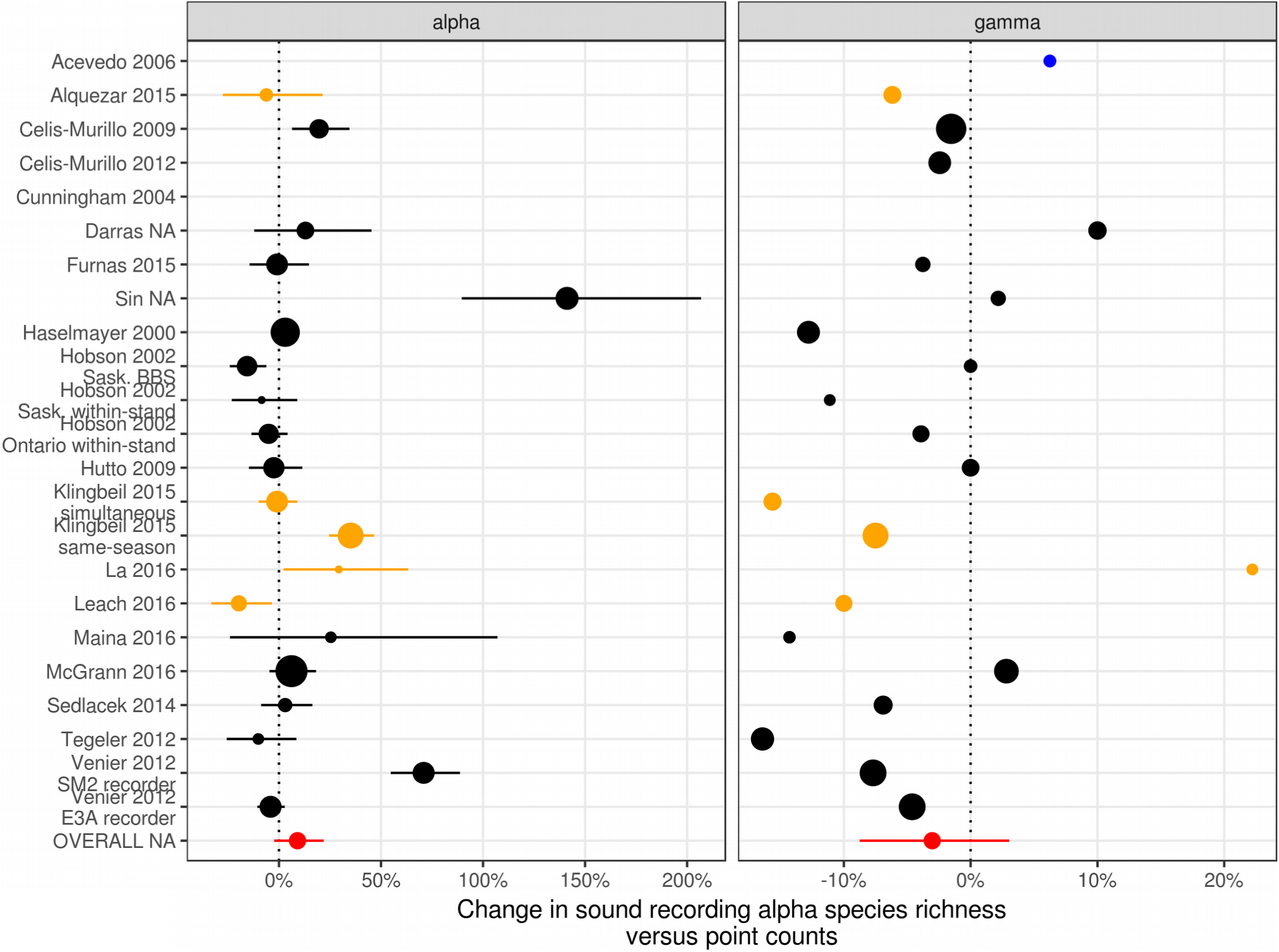
Response ratios of bird species richness sampled by automated sound recordings compared to point counts. The error bars display 95% confidence intervals, and indicate a significant (P<0.05) difference to the control (point counts) when they do not overlap the dotted line. The dot size is proportional to the sample size: sites number for alpha richness and total survey time for gamma richness. Orange dots represent studies in which, because of lack of detection distance data, detection ranges were greater for point counts than for sound recorders, while blue dots represent studies where the detection range was larger for sound recordings.

We found that microphone signal-to-noise ratio was a significant, positive predictor for the alpha (estimate: 0.086, P=0.03) and gamma richness log response ratios (estimate: 0.007, P=0.02) in the unlimited range scenario (Fig 2 and Table S1).

**Figure 2:**
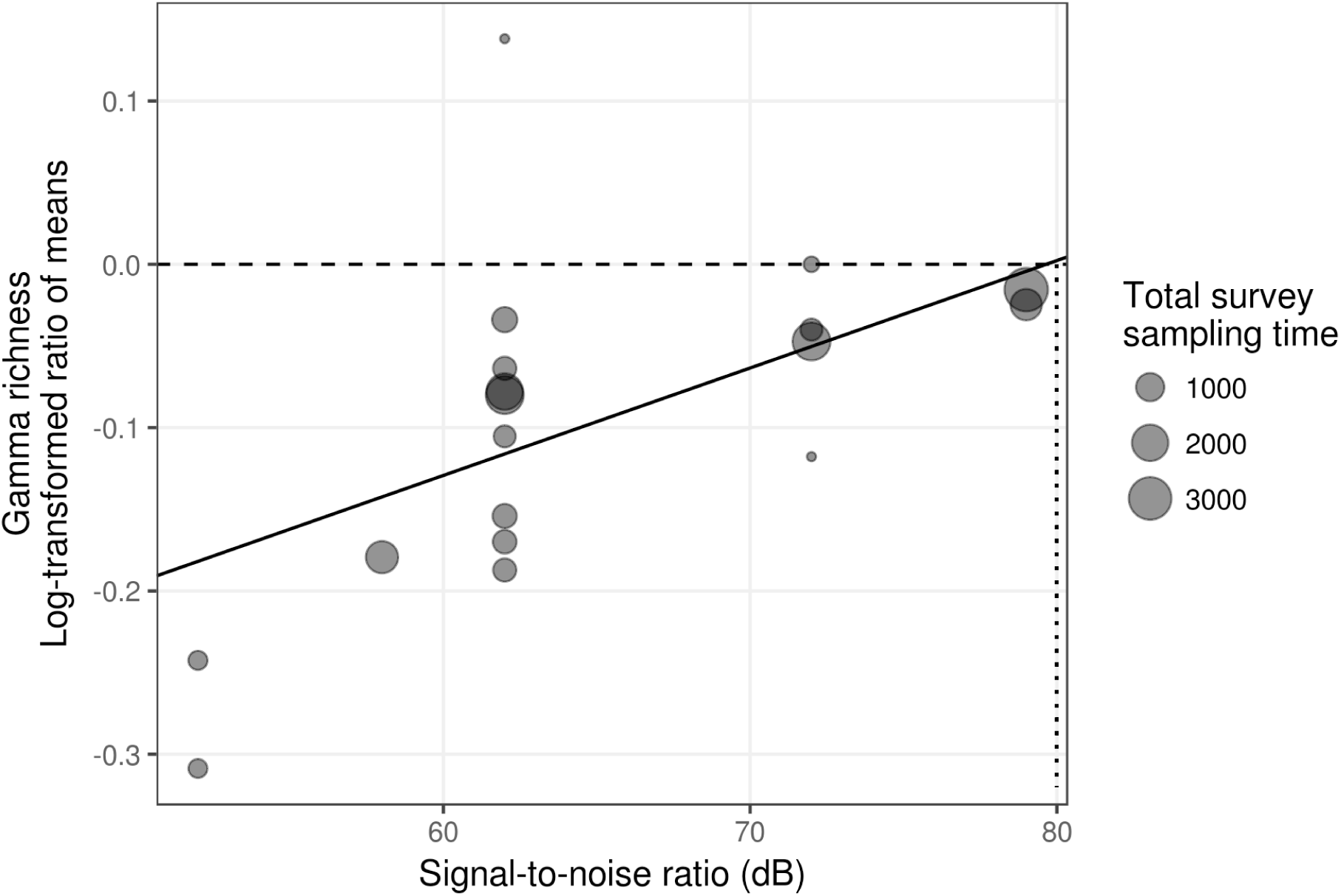
The relationship between microphone signal-to-noise ratio and log-transformed ratio of means for bird gamma richness. Positive values indicate higher gamma species richness in sound recordings versus point counts. The displayed regression line is based on a linear mixed effects model with signal-to-noise ratio as predictor, study ID as random effect, and total survey sampling time as weight. The line reaches zero at approximately 80 dB.

On average, species detected by both methods made up 81% of the total species count, both for identical and unlimited range detection areas. Respectively for the unlimited and identical range scenarios, 14% and 12% of all species were only detected with point counts, while 5% and 7% were only detected in sound recordings.

### Field studies

#### Bird abundance and richness in forest surveys

In our field survey, bird abundance and species richness did not differ between survey methods (Fig S2). Paired-sample Wilcoxon signed rank tests for comparing mean richness and abundance between sampling methods were statistically not significant (richness estimate: 1.50, *P* = 0.27; abundance estimate: 0.50, *P* = 0.16). Total richness reached 66 species in sound recordings, while only 58 were found in point counts.

#### Detection distance in forest surveys

Direct distances to detected birds were distributed differently between survey methods. Figure 3 shows how the detectability of both methods deviates from perfect conditions, where we would see an even histogram depicting equal detection rates in all area bins. Point counts showed a less steep decline in the number of detections with distance compared to sound recordings, but they also have a markedly lower detection rate at close distances.

**Figure 3:**
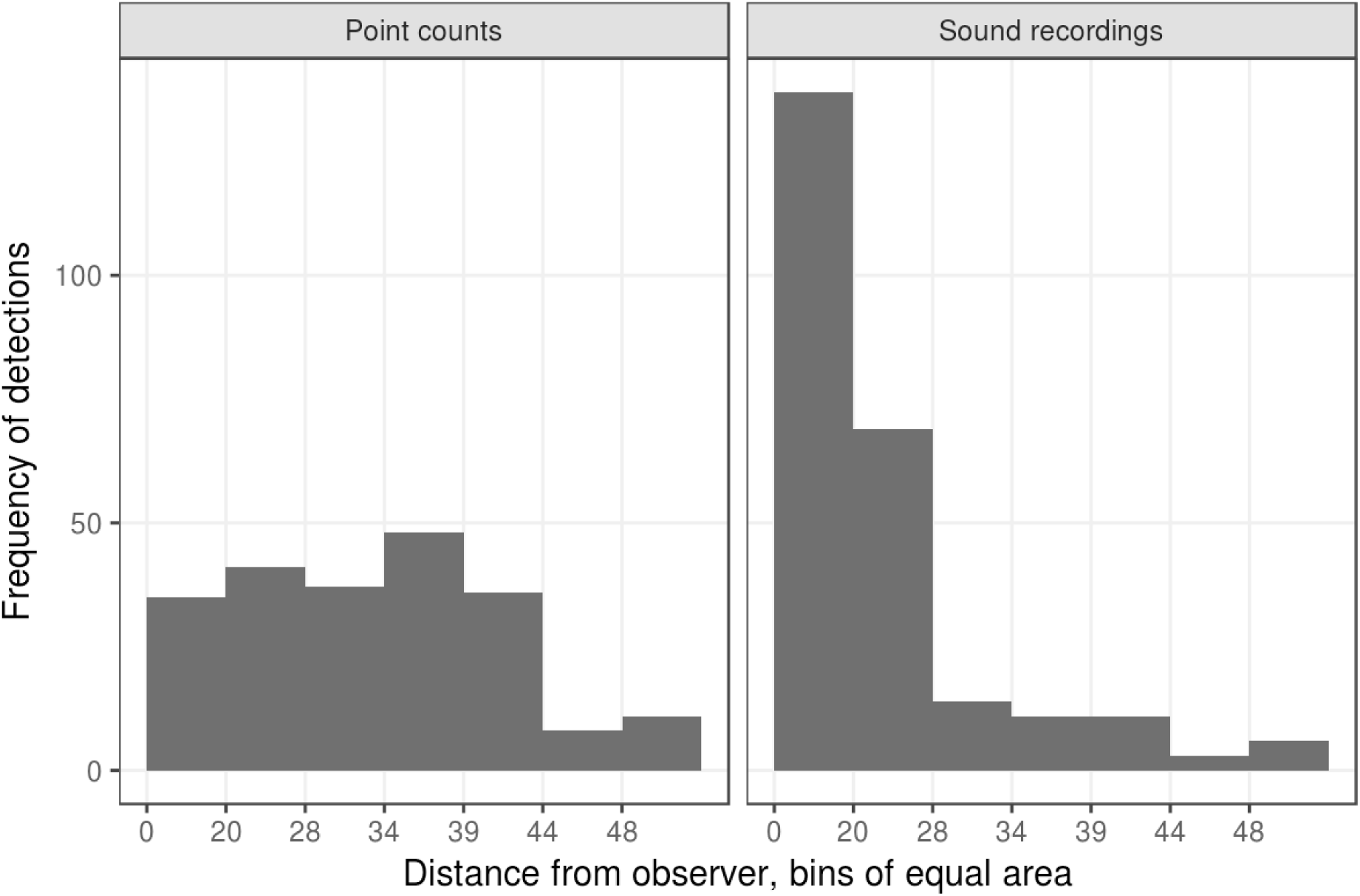
Frequency of bird detections with distance from the observer. Distances on the x-axis are separated in bins of equal area. Only 3% of detections are above 50 m.

#### Distance estimation accuracy

Estimated distance from sound recordings strongly correlated with the actual distances of the birds (Fig 4). The distances to the raucous call of Acridotheres javanicus correlated slightly less and were relatively over-estimated.

**Figure 4:**
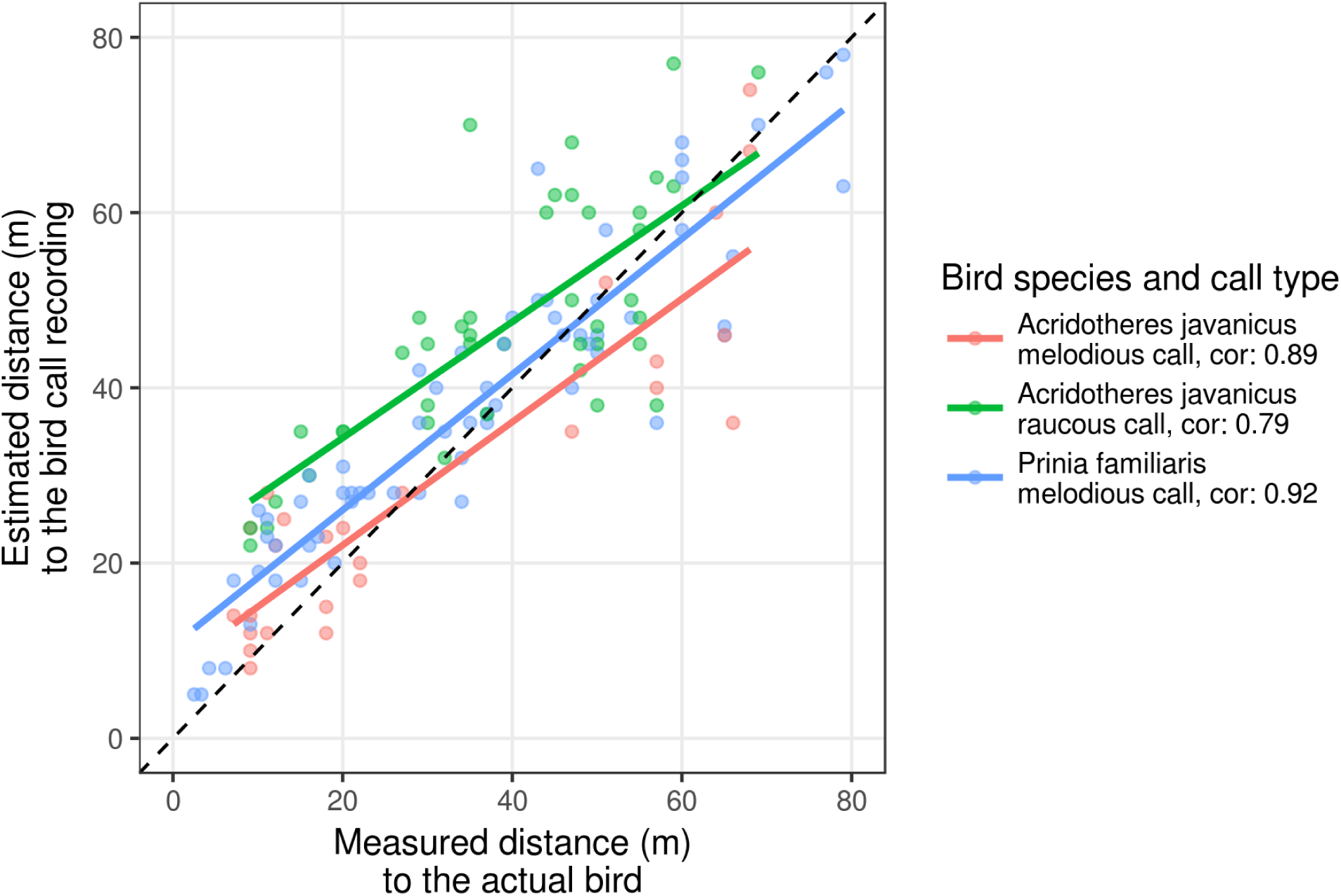
The relationship between distances estimated aurally from sound recordings and laser-measured distances to birds for two different bird species, one of which emitted two different call types

## Discussion

### Technical differences between humans and microphones

More often than not, people interested in autonomous sound recording systems ask: “What is the detection range of that system?”. The answer is complex, as sound detection spaces have many determinants, like habitat structure, ambient sound level, as well as source sound level, height, and directivity (Darras et al., 2016). The variables that ecologists have influence on however, are the technical specifications of the recording systems.

Species detectability in sound recordings depends foremost on the microphone’s sound quality. For our purposes, sound quality is best described by the signal-to-noise ratio, which indicates the microphone sound level output relative to its self-noise floor (usually measured with a 1 kHz, 94 dB SPL reference sound). The sound recorder electronic circuitry (pre-amplifiers) also add some noise geometrically to the microphone's self floor, but that figure is relatively small in comparison. In contrast, microphone sensitivity, which is more often reported than the signal-to-noise ratio, merely describes the ratio between the analog input (sound pressure) to the electrical output (analog or digital) of the microphone and indicates the purpose of the microphone. Microphones for soundscape recordings are usually chosen with a sensitivity that is adapted to record the relatively low sound levels of wildlife (in contrast to a microphone for rock music vocalists) without clipping of the output signal. The final sound output level can additionally be adjusted in the recorder by signal amplification (usually digital), and as long as the recorded audio is not clipped (ie. does not surpass the microphone’s output capacity), sound level can be corrected further by postprocessing on a computer. It follows that microphone sensitivity is usually quite irrelevant. Signal-to-noise ratio however, cannot be altered or enhanced, it is a constant characteristic of the microphone which tells how faithfully and cleanly a microphone records sound.

The higher the signal-to-noise ratio, the higher the quality of the recordings since there is less intrinsic noise added by the microphone itself. Thus, the higher the signal-to noise ratio, the better far away calls that would be correspondingly faint in sound recordings can be heard or enhanced (by digital amplification or analog increase of the loudspeakers’ volume) until they become audible. Thus increasing microphone signal-to-noise ratios improve detection and identification probability by extending their range. The only requirement is that the recorded sound levels of interest are above the microphone’s self-noise. But it also follows that in natural recording settings, the ultimate limit to a sound recording system’s detection range is the ambient sound level. Even the best microphones with signal-to-noise ratios below the ambient sound level cannot record far away bird sounds as soon as they drop below that point. Ambient sound level is made of “noise” sources like such as anthropogenic (i.e. engines), biophonic (i.e. insect noise), or geophonic (i.e. wind, rain) sounds, but also of all the other sounds of interest (like bird vocalizations).

Another important point is the fundamental difference between microphones and human ears in how they respond to different sound levels. Microphones are linear transducers of sound energy, which explains their limited dynamic range, as microphones are either optimized for loud or for faint sounds by their sensitivity level. The human auditory system, however, responds differently to sound energy: faint sounds are amplified, and loud sounds are dampened, resulting in the impressive dynamic range of our ears (Stevens and Warshofsky, 1981). The latter phenomenon explains why we have a steep fall-off in the number of bird detections in sound recordings with distance (Fig 4), which is also apparent in the reduced number of detections in simultaneous recordings (Fig S2), especially at greater distances. Thus in our study, our detection range is relatively limited compared to point counts, due to the weak signal-to-noise ratio of our microphones, and it likely similar in all the other studies which used the same microphones.

### Systematic review

The main challenge for comparing richness measures from the two different survey methods was to enable a fair comparison by using equal sampling areas for both methods. Most studies addressed that problem, but in several cases, that issue was not addressed, and sometimes distance data were not collected either to alleviate that problem. It resulted in comparisons between unequal sampling areas, usually biased towards point counts (Alquezar and Machado, 2015; Klingbeil and Willig, 2015; La and Nudds, 2016; Leach et al., 2016).

In our field study, we used modestly quiet microphones: one SMX-II microphone with the common Panasonic WM-61A element (signal-to-noise ratio >62 dB), with one Knowles SPM0404UD5 element (signal-to-noise ratio >59 dB, this element was used for ultrasound recordings at night). Most studies in this meta-analysis used similar equipment (Song Meters from Wildlife Acoustics) with the same microphone element. We estimated that in our forest sites, there were only 5% of the total detections above 45 m, indicating roughly the “range” that these microphones deliver. Hutto et al. (2009) showed that with the same microphone elements in their open survey habitats, bird vocalisations above 100 m are mainly undetected in sound recordings because they are too distant. Venier et al. (2012)also demonstrated that most bird calls above 50 m are undetected with the same microphone elements. Sedlacek et al. (2015) also suggest that the signal-to-noise ratio is so low for sounds above 50 m that detectability is significantly decreased. Richness values from studies using unlimited radius point count data, that showed the strongest bias towards point counts, were entirely reversed when the distance data were used to equalize detection ranges between methods (eg. Hutto and Stutzman, 2009, Sin et al. unpublished data).

More recent microphones used since 2015 by the same manufacturer are much more accurate (SMM-A2 microphones, signal-to-noise ratio 80dB) and compare well to the more expensive setups that were used by other authors in this review (ME-62 from Sennheiser and Compression Zone Microphone from River Forks Research Corporation). In their studies, the recording system was also evaluated and deemed to pick up far away sounds as well as human ears, necessitating no truncation of the corresponding point count detections data set. At the other extreme, cheap microphones with low signal-to-noise ratio were used in several studies and shown to have a maximum range of 40 m (Furnas and Callas, 2015; McGrann and Furnas, 2016, Sin et al. unpublished data) or estimated to have even lower detection ranges of 20 m (Maina et al., 2016, pers. comm.). These were still comparable with the point counts data when truncated to these respective distances.

All in all, this underlines that while human observers presumably have a consistent detection range, the detection range of sound recorders is certainly very variable: it is dictated by their microphones’ signal-to-noise ratio, which is a decision made in every study design, constrained for budgetary reasons. The dramatic influence of the signal-to-noise ratio was demonstrated in our analysis of the log response ratios in the unlimited range scenario (Fig 2), from which we can infer that higher-end microphones can be on par with human hearing by sampling sound detection spaces as large as those sampled by human observers.

Compared to point counts, the methodology of sound recording surveys itself is not standardized. Point counts have been performed for decades, and are most often based on a variation of the survey design described by Ralph et al. (1998). Sound recording setups however, differ with respect to the microphones’ placement, which affects their detection range greatly. Installation of microphones on the ground places them in sound shadows (Morton 1975), effectively hiding them from the sound field. The number of microphones is also critical: choosing a single audio channel (mono configuration) leads to a loss of auditory and spatial information, which is necessary for estimating bird distances accurately. In the worst recording setup, placing low-cost (low signal-to-noise ratio) microphones on the ground while recording only one channel is equivalent to conducting a point count with a hearing-impaired person lying on the ground while closing one ear. Thus, many of the presently reviewed studies suffered from a tactical disadvantage from the outset.

A further argument in favor of sound recorders is that almost all methodological comparison studies discussed here disadvantaged sound recorders because identical time intervals were compared, whereas autonomous sound recording units are capable of sampling continuously or at random times distributed through the day or the season. The advantage of autonomous sound recording units, when used with such a monitoring program that makes better use of them, is impressively demonstrated by Klingbeil et al. (2015) who showed a 35 % increase in alpha diversity (Table 1), although gamma diversity admittedly stayed slightly inferior, probably due to the smaller detection range of their sound recorders compared to point counts. Furnas and Callas (2015) also demonstrated how detection probability varies for different species with time of the day, also supporting the idea that sampling times distributed throughout the day will sample the entire bird community much more effectively, which is also echoed in La et al. (2016) where the most efficient sampling schedule for detecting the entire bird community was tested.

### Meta-analysis

We chose to present two survey method comparisons in our meta-analysis: one with identical detection ranges, and the other with unrestricted/unlimited detection ranges. Both account for differences in sampling area in different ways. While the identical range scenario simply truncates data sets to a common detection distance and allows the direct comparison of mean richness values, the unrestricted range scenario accounts for differences in sampling area by including co-variates in the statistical model: out of microphone number, height, and signal-to-noise ratio, we found that the latter had the highest predictive power and used it to account for the different effective sampling areas. It was found that signal-to-noise ratio was a significant and strong positive predictor of the sampling success of sound recording systems. Both meta-analyses thus pointed to the same conclusion in different ways, namely that sampled species richness numbers do not differ between both survey methods when sampling areas are equal, and that in sound recordings, the sound detection area increases with signal-to-noise ratio, up to a point where both methods are perform equally.

When we examine the underlying detection rate profiles for each survey method more in detail (Fig 4) however, it appears that even with equal detection ranges, point counts and sound recorders do not sample the same area identically. Point counts often introduce avoidance zones around the center (eg. Supp. Mat. in Prabowo et al., 2016) and keep higher detection rates with increasing distance, while sound recording detection rates are focused on the center and rapidly fall off with distance. This points to the fact that distance sampling should be used to account for these different patterns to estimate abundance or density of birds. One should account for the different detectabilities of loud and quiet birds to compute richness estimates also. However distance estimation methods from sound recordings are not established yet, and estimated distance data was only available in our field study because of our particular distance estimation protocol using supporting sound transmission sequences. Therefore, when distance sampling is not used, it is preferable to truncate detection data sets to a common threshold distance below which there are no detectability differences between the treatments of interest (Schieck, 1997), an approach that we put forward in our meta-analysis with the identical detection ranges strategy.

### Comparison of bird point counts versus automated sound recordings

<u>Table 2</u>: comparison of advantages and disadvantages of human observation (point count) and automated sound recording methods for surveying birds.

#### a) Standardization and verifiability

Point counts suffer from a trade-off between observation time and sampler bias: with an increasing number of observers more simultaneous - and thus temporally unbiased - data points can be obtained, but the number of observer-specific - thus sampler biased - data points increases. The considerable magnitude of the observer bias has been convincingly demonstrated by Alldredge et al. (Alldredge et al., 2007a) and Simons et al. (2007). In contrast, sound recorders incur no sampler bias (or recorder bias), provided that microphones are calibrated. Microphones are manufactured within given signal-to-noise ratio tolerances to start with, but signal-to-noise ratio may drift apart with time, depending on the environmental stress they have experienced (rainfall, temperature variations, mechanical shocks, etc.). Thus, regular measurement of microphone signal-to-noise ratio at different frequencies are required to ensure that they can be calibrated, so that different recording units have the same detection efficiency. Alternatively, reference recordings such as our sound transmission sequence can help to gauge bird call distances in recordings. Even so, uncalibrated microphones only result in different detection probabilities, and do not incur biases in bird identification like different human observers do.

Verifiability is low for point count surveys as we are essentially depending on the identification skills, current physical state, and memory of a single observer. Especially in tropical regions, the higher number of species vocalizing simultaneously makes correct identification of all individuals a challenging task. Moreover, auditory detections are sometimes uncertain (Mortimer and Greene, 2017). When point count observations have corresponding photographic or audio evidence material, the bias between observers can be lessened, but these verification data are seldom available. The bias can also be corrected by taking other preventive measures (Lindenmayer et al., 2009). To obtain verification data however, an additional worker is usually needed, further raising the costs of the survey. With sound recordings, audio material is essentially available at no additional cost, and it can be used at leisure for future identification checks. Venier et al (2012) showed how field survey data can be corrected using sound recordings and how greater species counts can be achieved after re-interpreting them. Even if sound recordings were processed by people with different or no experience, as long as the same expert ornithologist reviews all observations, the species identification can be standardized, which is particularly helpful in long-term monitoring projects.

#### b) Sampling completeness

Several factors affect the sampling completeness, one of which is the avoidance effect introduced by human observers. Shy species are usually affected by the presence of human observers, especially when there is more than one (Hutto and Mosconi, 1981). Disturbance effects from observers on birds are not well documented (but see Fernández-Juricic et al., 2001). On the contrary, it is also possible that some birds are attracted by human presence. In our field study, birds are relatively less often detected close to observer (Fig 4), where detectability should be highest. In contrast, bird detections in sound recordings are most often near to the recorder. We interpret these diverging patterns as evidence of a net avoidance effect in point counts, due to disturbing human presence. At best, this avoidance effect diminishes bird detectability at close range if disturbed birds simply move further away while still being detected, but at worst, intolerant birds would be deterred and leave the detection area of the observer. Note however that this avoidance effect is dependent on the bird community (more secretive birds versus birds adapted to anthropogenic disturbance), and as Prabowo et al. (2016,supplementary materials) illustrated, birds in disturbed systems may exhibit that trend to a lesser degree. We do not discuss the lower number of detections in sound recordings at large distances since this was shown to be mainly a function of the microphones' signal-to-noise ratio.

Point count data include visual detections too, which is an undeniable advantage of point counts, especially in open habitats, where visual detections are more common. However, even survey comparisons in open habitats do not yield a large advantage for point counts (Alquezar and Machado, 2015; Celis-Murillo et al., 2012), and they do not necessarily have considerably more visual observations. In our study, only 5% of observations in our dataset were visual only, against 1% in Furnas and Callas (2015). Also, given enough (recording) time, most birds vocalize eventually by song or by call, so that most birds can be detected using passive acoustic monitoring. Finally, we note that in simultaneous surveys using both methods, the average percentage of species detected by both methods was relatively high (81%), with only 12% of species detected only in point counts.

In point counts of species-rich sites, birds can be missed when they occur simultaneously or because of human error (fatigue and lack of attention), especially during dawn chorus or during the first minutes of a point count (Hutto and Stutzman, 2009). Hutto et al. (2009) also showed that two thirds of the birds that were detected by autonomous recording units but failed to be detected in point counts were simply overlooked in field. Also abundance numbers can be underestimated for abundant birds (Bart and Schoultz, 1984). In contrast, sound recordings can be played back repeatedly, often leading to tangible advantages for infrequently vocalizing birds (Celis-Murillo et al., 2012), so that birds can only be missed if the listening time is constrained artificially or for budgetary reasons. Furthermore, spectrograms (e.g. sonograms) are routinely generated and inspected, while listening to audio recordings, so that bird calls are detected both visually and aurally, further enhancing detection probability compared to the aural-only detection of birds in point counts.

There is some debate as to whether sound recordings are more or less effective than point counts for detecting rare birds. As Celis Murillo et al. (2012) pointed out, some authors found that point counts were more effective in some studies (Haselmayer and Quinn, 2000; Hutto and Stutzman, 2009). One possible reason is that since visual cues are available, rare birds can be identified with more certainty (Hutto and Stutzman, 2009; Leach et al., 2016), however in both of these studies, the sound recording systems had smaller detection ranges than the unlimited radius point counts they were compared to. Venier et al. (2012) on the other hand, argue that detecting rare species is more cost-effective in extensive data sets collected by sound recorders. Other than that, for vocalizing birds and at identical detection ranges, there is no reason why rare birds should be inherently more easily detectable with one or the other method.

Consequently, since detection probability is a function of sampling intensity, and sampling intensity is much more easily extended with sound recordings, passive acoustic monitoring systems have a much greater potential for detecting rare species or confidently concluding their absence (Tegeler et al., 2012), especially when combined with automated identification algorithms.

#### c) Data types and compatibility

##### Data types

When visual detections are numerous, human observations can yield supplementary data about behavior (nest defence, territorial fights), consumed food items, stratum of occurrence in the vegetation, sometimes even the sex and age of the bird, although such data are rarely used and it is challenging to get a meaningful dataset that is suitable for statistical analyses. To some degree, bird calls also convey information that is accessible in sound recordings, since calls have different functions: territorial advertisement, mate attraction, and alarm calls all give information about the bird’s current behavior. Also, distinguishing between songs - which are typically territorial - and calls can give cues about whether the habitat is suitable for breeding or is rather only visited by stray birds. In some cases when the bird is moving and calling at the same time, movement can also be reconstructed, especially when using microphone arrays (Blumstein et al., 2011). Depending on the position of the microphones, it is also possible to reliably pinpoint the animals’ position in space (Bower and Clark, 2005) and deduce the calling height and stratum of occurrence.

Sound recordings provide continuous sound recordings where human observation can only provide a filtered sample of the original visual and aural observation. It enables to derive secondary data types that are not accessible with point count data. For example it is much easier to measure the call activity in time units or call rates, which can be an interesting alternative to abundance measurements. This relates more accurately with bird activity, which is a more relevant measure for functional analyses. Cunningham et al. (2004) showed that although vocal activity and bird abundance are significantly related, they are only weakly so, meaning that calling activity may represent an alternative to abundance measures, rather than a proxy for it. With continuous sound recordings, we can also analyse temporal dynamics throughout the day, between days and between seasons and assess phenological trends and temporal dynamics (Lellouch et al., 2014). Temporal beta diversity and species turnover can be analysed at any temporal scale and it is a major factor influencing species richness (Flohre et al., 2011), when alpha diversity can even be misleading (Tylianakis et al., 2005). Furthermore, we can generate sound diversity indices for large datasets automatically, at the only expense of coding and computation time. Sound diversity indices have been used repeatedly and usually correlate well with field measures of species richness (Depraetere et al., 2012). Finally, all other sonant animal taxa are also available, allowing a more holistic biodiversity survey.

#### d) Comparability and compatibility

Two essential data types that are derived both from human observation and sound recording data are abundance and distance data. Both are obtained in different ways and it is worth to examine them in more detail.

Abundance data tend to be more readily obtained from point counts, since it is more intuitive to estimate the position of the birds and relate it to previous activity as to guess the bird abundance on the sampling site. However, especially in dense habitats, bird individuals are rarely seen and thus hard to distinguish so that we can never really know for sure whether two different non-simultaneous sightings correspond to the same or different individuals. A more conservative estimate of abundance is the maximal number of simultaneously detected individuals of one species. It has been used before as a conservative measure of abundance (Clough et al., 2016; Teuscher et al., 2015), which was also our measure of choice. It can be obtained from both survey methods and allows to compare or merge datasets while using an identical measure of abundance. Still, it is also possible to count uniquely identified individuals in stereo recordings in a similarly inuitive manner as in point counts (Hobson et al., 2002), because the birds’ location is audible. Similarly, individual birds have unique calls which can be distinguished from another upon close analysis (Beer, 1971). Simple presence/absence data can also be merged without compatibility issues, as is common in occupancy studies (McGrann and Furnas, 2016), and acoustic data sets have been merged with point count data in several other recent studies (eg.Leach et al., 2016; McGrann and Furnas, 2016). Sedlacek et al. (2015) also computed abundance values based on acoustic recordings. Finally, sound recordings may be more amenable to calculate bird density when they are used to sample audio snapshots several times a day (Buckland and Handel, 2006).

The estimation of bird distances in point counts can be challenging and inaccurate (Alldredge et al., 2007b). We must bear in mind, however, that even though the distance is measured, it is also an estimation because it is based on the estimated position of the animal, except in the few cases when it can be seen. In the more challenging case when the animal is moving, both sound recordings and point counts have similar issues because distances should actually be measured multiple times, or a rule be set for determining which distance to choose among the many recorded distances. We have shown in our experiment that distances can be estimated accurately with the help of reference material like our sound transmission sequence. One should note that atypical calls can be more challenging: in our experiment the raucous vocalizations of *Acridotheres javanicus* is more variable in intensity than typical broadcast calls, explaining the slight over estimation of their distances. However, we still attained a high correlation between measured and estimated distances. Presumably, the harmonic frequencies of bird vocalisations or lack thereof, which are easily visualised in spectrograms, serve as cues for the estimation of the distance. Previously, Hobson et al. (2002) suggested that spectrograms of bird calls can also be used to estimate their distance with reference to their sound source level. When microphones are calibrated and transmission patterns and source call sound levels are known, it is possible to calculate a distance, however that approach would be more tedious than the one we used. Nevertheless, the accuracy of distance estimations from sound recordings should be evaluated in more detail with a variety of estimators, land use systems and directions, and if needed, new, more accurate methods should be devised.

#### e) Practicality

##### Travel time

Observers carrying out point counts usually only need to reach the sampling site once before the survey to become familiar with the itinerary and surroundings. For every subsequent data collection, only one travel is necessary. In contrast, sound recorders need to be installed before they start recording and must be picked up for collecting the data or recharging batteries (but see Aide et al., 2013 for an example of remote data collection and continuous power supply). Depending on the study design, either one of the survey methods could be more practical: if sampling rounds on consecutive days at the same site are needed and the theft risk is low, sound recorders will prove handy. If the number of sampling sites is high and replicate visits are few, either many recorders or frequent travels will be needed.

##### Scalability

Due to the limited number of observers, the survey time for point count surveys needs to be optimised so that all target sites can be reached and enough time is spent on each site. Acoustic surveys, however, allow for greater flexibility in scaling up sampling effort, provided enough recorders are available. It is effortless to program automated sound recorders to record for more hours, or even the whole day, which only comes at the expense of data storage and processing time, as well as energy supply, both of which are generally cheap when compared with specialised labour (i.e. ornithologists). It must also be stressed that in species occupancy modeling, the number of replicate surveys considerably improves site-level detectability, overall accuracy and precision of state variables such as richness. It follows that autonomous recording units confer a considerable advantage in that respect because of the ease with which replicate surveys can be conducted.

##### Expert workforce

It is often costly to hire taxonomic experts for traditional field surveys, which require their presence. Passive acoustic monitoring systems, however, can be installed and picked up by ornithologically inexperienced staff, whereas the financial allocation for taxonomic experts can be minimized to use them only for the proper identification of the animals, or even postponed for the time when funds become available. Moreover, since their presence is not needed, data can be sent to them or accessed online (see http://soundefforts.uni-goettingen.de/), helping to keep travel costs low. It is often stated that identifying birds inside sound recordings is a time-consuming process, but solutions exist for shortening the processing time (Zhang et al., 2015), and it is always possible to restrict the listening time strictly to the time of the length of one broadcast, thereby simulating a “blind” point count.

In the near future, automated species identification is also conceivable so that reliance on expert ornithologists will be even further diminished. Numerous studies have showed how calls of single species can be detected with a measurable probability and accuracy using computer algorithms (e.g. Brandes, 2008). Tegeler et al. (2012) demonstrated how automated analysis of sound recordings could significantly increase the total detected species count. The number of species that can be reliably identified computationally will unrelentingly increase, but it remains to be seen whether more complex song structures and entire song repertoires can be reliably assigned to species. Considerable input from human experts will be needed even with the most “intelligent” automated methods like machine learning. As for automatic identification of bats, some commercial solutions already exist for bats from temperate regions (Wildlife acoustics), but tropical bat communities are much more challenging.

##### Material costs

In the case of birds, point counts usually only require binoculars and field gear. However, it is often the case that birders use their own, helping to keep the costs down. Autonomous sound recorders are however mostly quite costly, although a multitude of hardware solutions exist (see Merchant et al., 2015 for an overview), from do-it-yourself constructions (e.g. Maina et al., 2016; Whytock and Christie, 2016) to commercial products (e.g. Wildlife acoustics) spanning a price range of hundreds to thousands of USD. An important consideration is also that in longterm studies, it is often difficult to hire the same people throughout, and ornithologists are rarely hired for conducting field surveys full-time. Sound recorders, however, are pieces of hardware that are purchased once and typically last for years if maintained properly. They can be used over and over again until irreparably broken or stolen, greatly facilitating long-term data compatibility. Then again, commercially available recorders have limited life cycles and their microphones are eventually discontinued. The availability and specifications of such products are dictated by economic and marketing considerations, so that it is problematic to guarantee constant audio specifications of passive acoustic monitoring systems for decades.

##### Transportability and usability

Some wilderness habitats in forest, at high elevations, or unexplored regions can be very difficult to reach. Especially when conducting morning point counts of birds, the observer preferably has to be present on-site at dawn. This is often impossible in inaccessible areas where travel by night would be required, or dangerous in sensitive areas. When using sound recording platforms, however, as long as the recorder is installed before its programmed recording time, we can always reliably and exactly meet the desired schedule.

Prevost (2016) showed that acoustic recording systems, due to their low size and weight, were amenable to installation on air balloons to sample from elevated points of sound. Also deployment of sound recording units to inaccessible acreas with unmanned aerial vehicles should be feasible. In the future, large geographical scales could also be sampled using autonomous wireless sensor networks (Collins et al., 2006).

Autonomous recording units are easy to use in the field and the lab. Sound recorders can be set up as quickly as humans need time for getting ready for the point count. In our own study, with pre-charged and pre-programmed commercial devices, installation can be completed in two minutes. With some less practical designs, installation can require half an hour (Hutto and Stutzman, 2009). However, although recorders can be set up rapidly, they suffer from a drawback when it is raining:, many microphones are not weather- or waterproof and when protective foams are used, they are usually soaked with water after rain, which results in a loss of sensitivity and can take between less than one hour or several hours to dry, depending on ambient humidity, based on our own experience with Song Meters (Wildlife Acoustics). This is a technical challenge waiting for a solution. The scheduling of sound recorders does not usually require programming experience (eg. Song Meters of Wildlife Acoustics), but some custom open-source solutions (eg. Solo recorder, see Whytock and Christie, 2016) need some form of command line tool knowledge.

##### Rapid assessments

Sound recordings could be used to rapidly assess the avian diversity of a sampling site. Alternatively, bird call types could be counted as a proxy for morphospecies, and audio recordings are particularly amenable to rapid visual screening as they can be represented as spectrograms. However, some birds have a large repertoire of song so they might bias that measure, and other birds are also mimicking other species. Still, on a more abstract level, sound diversity indices can be computed from soundscape recordings, providing an even faster measure of biodiversity (Sueur et al., 2008).

##### Spatial scale of recordings

Sound recorders have more detections in their central installation point because of inherent technical characteristics of microphones (Fig 4). This is an additional argument for sound recordings when bird surveys need to be carried out on small plots (homegardens, smallholdings, etc.) where human presence would affect birds in the entire plot, or even in open habitats, where human recorders are visible from far away. In sound recordings, the fact that the detection distance profile puts more weight on the center of the survey point is also useful, because environmental co-variates are usually measured close to the center, eventually enabling a closer spatial linkage between them and bird community variables.

To conclude, ecology is a field of constant improvement, and most of it is technological. We have shown that sound recording devices can detect species richness levels that are on par with human observers, and if used properly, they can surpass richness levels from point counts. Autonomous sound recording units are more practical, scalable, consistent, and provide verifiable results. There is even more room for improvements as automatic identification algorithms and sound archiving software solutions are developed, which will facilitate handling the huge amount of data that they are generating.

Even so, at the time of writing, machines do not replace humans yet quite. All audio data are ultimately identified, checked and reviewed by experts - ornithologists in the case of bird surveys - and the latter still fulfil an indispensable function. Some might fear that sound recording devices are putting ornithologists out of a job, however one might also argue the other way around: as bird survey data collection becomes accessible even to nonornithologists, the amount of gathered data will probably increase so much that demand for expert workforce could increase. Ornithologists will have to become more flexible as data can be analysed at any time, outside of survey seasons, from anywhere as long as the data can be downloaded. Thus of course, ornithologists will not become redundant; machines are not thinking yet, and even machine learning approaches require human operators. Bird-watching also still constitutes a major pastime for millions of people around the world, generating huge amounts of data likewise (eBird). Moreover, we will always need humans to observe birds, assess their behavior, study how they are ingrained in their complex ecosystems, and comprehend their role in the bigger picture.

## Acknowledgements

We declare that there are no conflicts of interest. We warmly thank all collaborators who shared data and provided additional explanations to carry out our meta-analysis. We thank Edho Walesa Prabowo for carrying out some of the forest bird surveys. We thank the following persons and organizations for granting us access to and use of their properties: village leaders and local plot owners.

## Data archiving

Data will be fully provided in the supplementary materials

## Supplementary materials

**Table S1:**
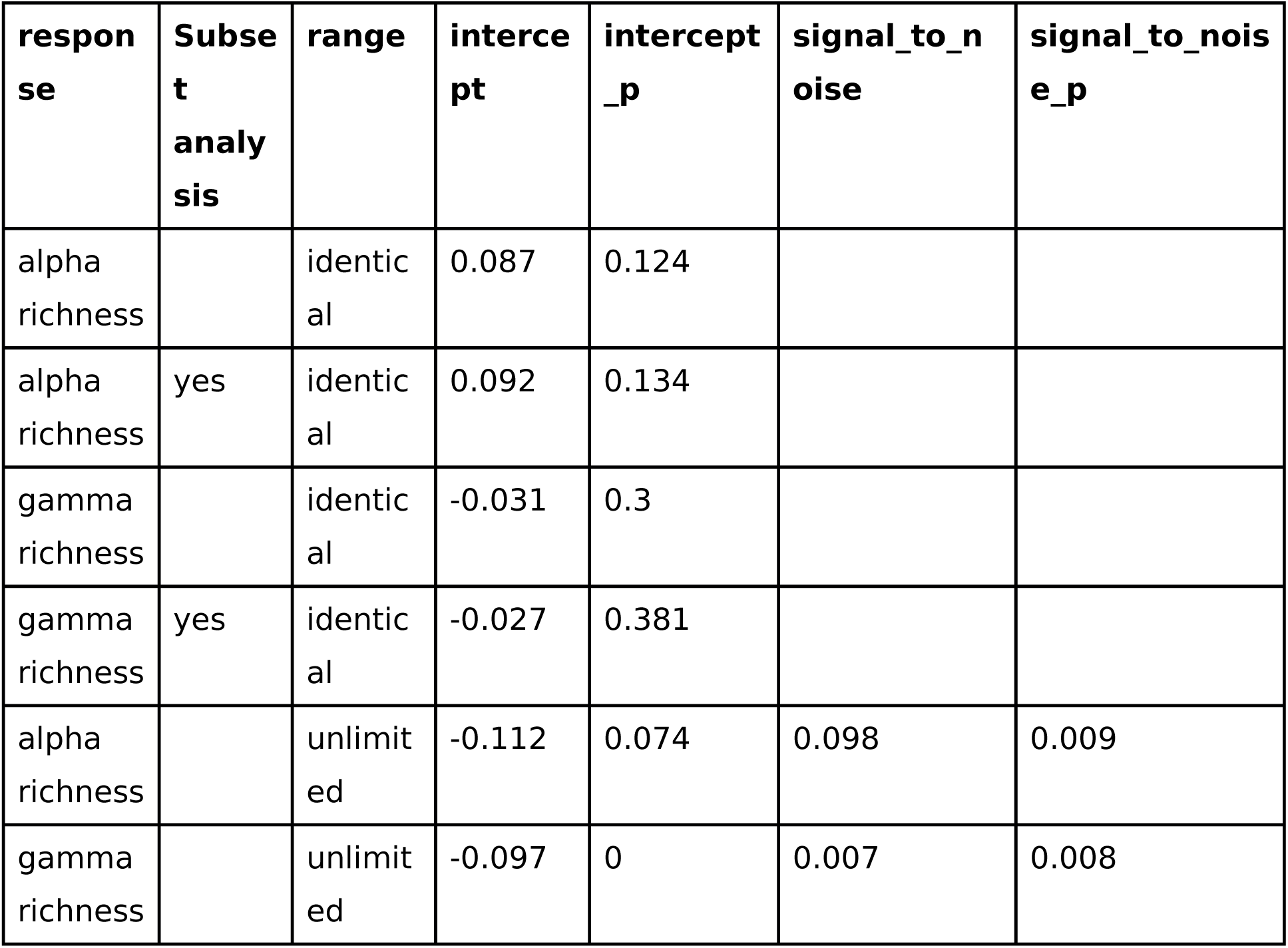
Multivariate meta-analysis and generalised mixed-effects model results for predicting alpha and gamma richness

**Figure S1:**
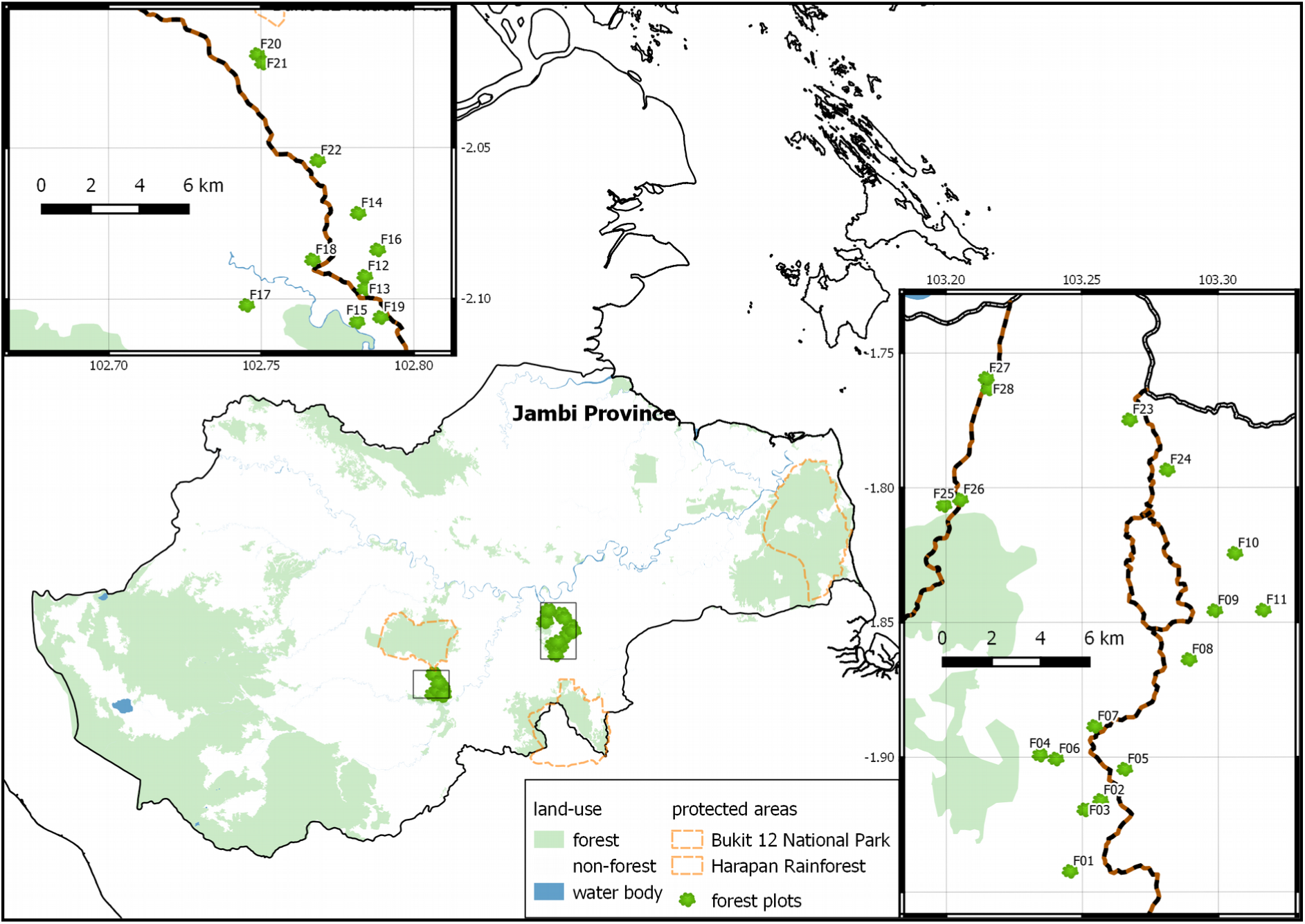
Map of the forest plots used in the survey

**Figure S2:**
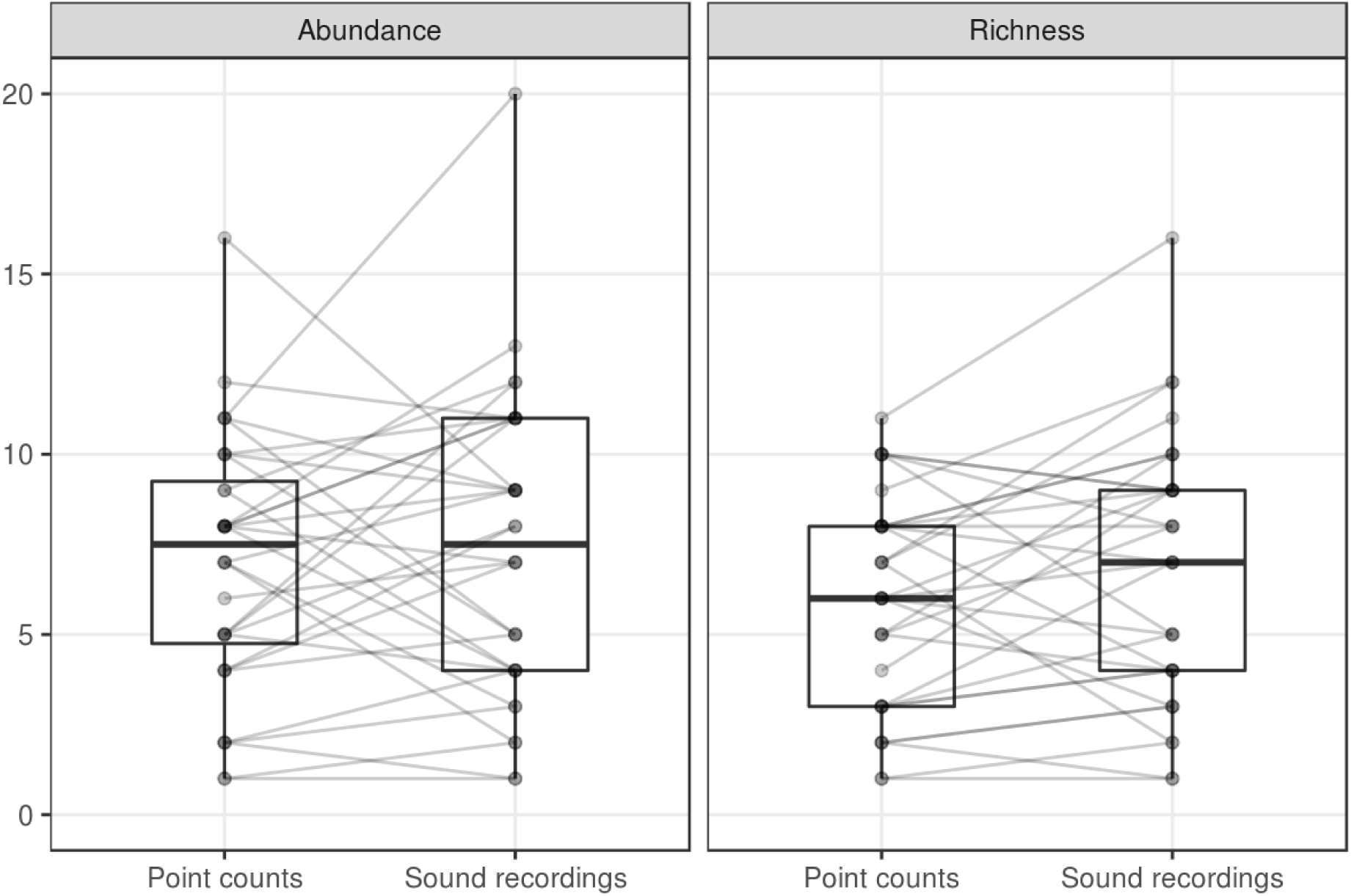
Bird richness and abundance sampled in 28 forest plots with 34 point counts and sound recordings, including all birds within 45 m from the sampler. Lines connect data points from identical plots, dots are partly transparent.

**Figure S3:**
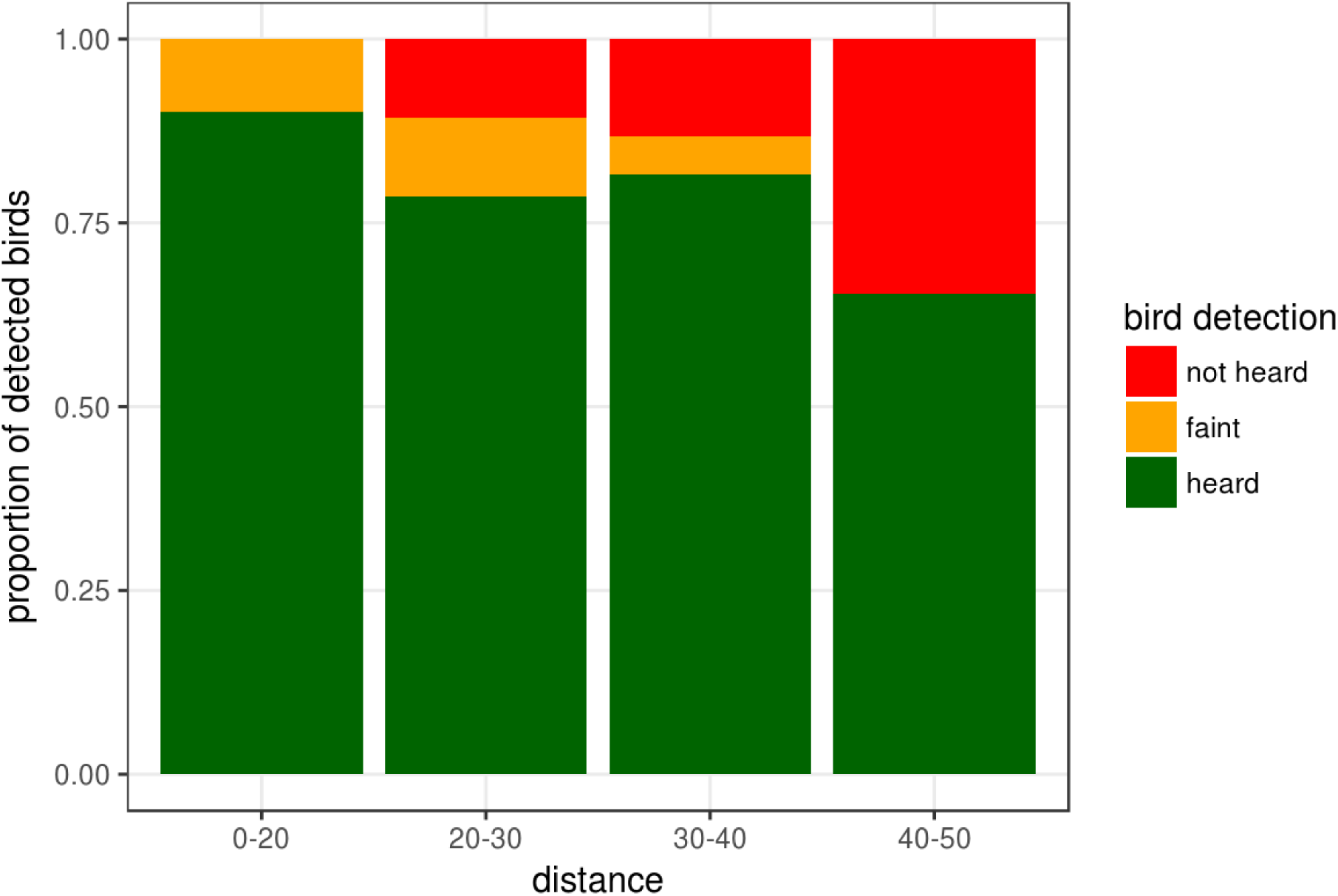
Proportion of birds detected in different distance classes in sound recordings that were concurrent with point counts.

**Figure S4:**
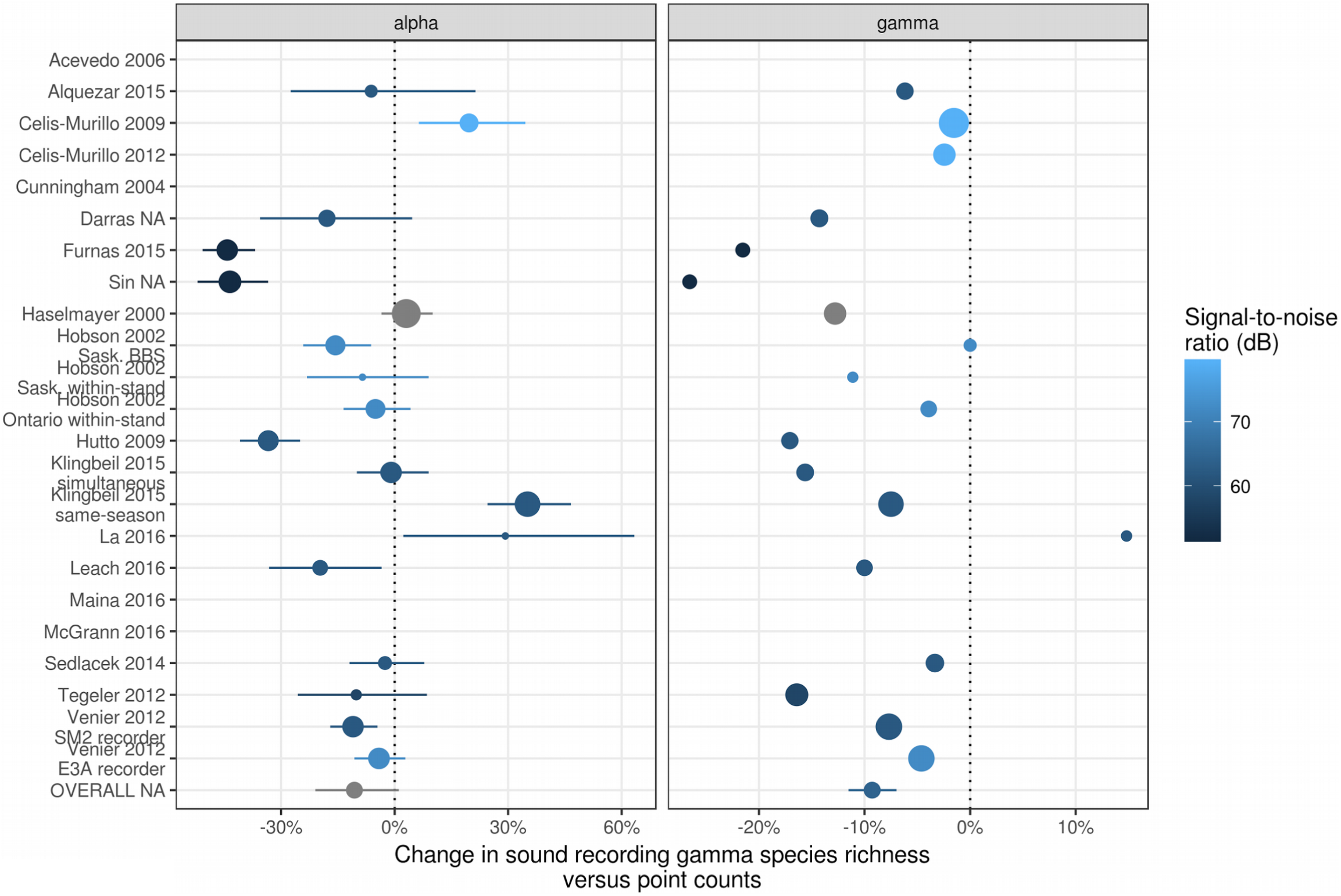
Response ratios for the unlimited range scenario. The overall value represents the response ratio at a species richness corresponding to an average signal-to-noise ratio.

